# Investigating insecticide susceptibility status of adult mosquitoes against some class of insecticides in Osogbo metropolis, Osun State, Nigeria

**DOI:** 10.1101/2023.02.16.528853

**Authors:** H. O. Raheem, Z. O. Iwalewa, L. O. Busari, K. A. Fasasi, M. A. Adeleke

## Abstract

The study evaluates the resistance and susceptibility of adult female *Anopheles gambiae* s. l., *Aedes aegypti* and *Culex quinquefasciatus* mosquitoes sourced within Osogbo metropolis, Osun State, Nigeria to four groups of insecticides [Permethrin, Deltamethrin, Pirimiphos-methyl and DDT (Dichlorodiphenyltrichloroethan)] and the distribution of their larval habitat within the metropolis. Mosquito larvae of the three genera were collected during the wet season and reared to adult stage in the laboratory. Emerged adult female mosquitoes were exposed to insecticide impregnated papers of the four insecticide groups for 60mins using WHO kits to determine the knock down rate (kdr). Thereafter, they were to holding tubes and left for 24hrs to assess their resistance and susceptibility according to the WHO protocol. Four types of larval habitats were identified (tires, ground pools, gutters and plastic containers). *Anopheles gambiae* s. l. showed the greatest resistance to Permethrin (49%) (p=0.04, p<0.05) while the highest susceptibility was recorded with Pirimiphos-methyl (69%) with the lowest against Permethrin (16%) (P=0.002; p<0.05). The greatest resistance of *A. aegypti* was against OC-Control (45%) (p=0.031; p<0.05). Permethrin had the highest susceptibility (60%) against *A. aegypti* while OC-control had the lowest (11%) (p= 0.005; p< 0.05). *Culex quinquefasciatus* had a lesser resistance to OC-control (38%) as compared with *Aedes aegypti* (45%). However, it was most least susceptible to Pirimiphos-methyl (52%) and DDT (17%) respectively (p=0.013; p<0.05). The susceptibility of *A. gambiae* s. l. and *C. quinquefasciatus* to Pirimiphos-methyl and *A. aegypti* to Permethrin is an indication of the possibility of success if employed for vector control of *A. gambiae* s. l., *C. quinquefasciatus* and *A. aegypti* respectively. This could be through their inclusion as active ingredients in insecticide treated nets (ITNs) and indoor residual spray (IRS) with a view to abating malaria and other life-threatening mosquito-borne diseases constituting global public
health scourge.

## Introduction

The public health menace of adult mosquitoes as vectors in the transmission of mosquito-borne diseases worldwide is worrisome. They transmit life-threatening diseases such as dengue fever, malaria, bancroftian filariasis, yellow fever, chikunguya fever, etc. (Grigoraki *et al.*, 2016). Mosquitoes of the genera *A. gambiae* s. l., *A. aegypti* and *C. quinquefasciatus* are most vectors responsible for the transmission of the deadly mosquito-borne diseases in human causing nearly a million deaths and over 700 million infections globally annually (Caraballo and King, 2014). Mosquitoes are cosmopolitan and are found in a variety of habitats including sewage water, stagnant water and fresh water (WHO, 2013). *Anopheles gambiae* s. l. are the vectors of malaria accounting for about 214 million cases of malaria leading to 438,000 deaths annually globally (WHO, 2016). About 1.1 billion people are at risk of contracting lymphatic filariasis (WHO, 2016), which is majorly transmitted by *C. quinquefasciatus* and species of *Anopheles* and *Mansonia. Culex tritaeniorhynchus* is the major vector of Japanese encephalitis, which is found in tropical and sub-tropical countries (CDC, 2016). *Aedes aegypti* and *A. albopictus* mosquitoes are mainly responsible for transmitting dengue and dengue hemorrhagic fever, yellow fever, and chikungunya. It was reported that dengue infections affect about 2.4 million persons annually (WHO, 2016).

Despite global efforts in the prevention of mosquito-borne diseases through preventive chemotherapy and vector control programmes, the burden of the diseases persists through the emergence of resistant vectors to insecticides used for vector control. Although, despite the six classes of insecticides namely, Organochlorines, Organophosphates, Carbamates, Pyrethroids, Pyrroles and Phenyl-pyrazoles insecticides are currently used in mosquito control programmes worldwide (WHO, 2016), resistance has still been extensively reported. Thus, the necessity for rapt understanding of the resistant mechanism adopted by mosquitoes with a view to making broad spectrum insecticides that will be employed in vector control programmes.

Nigeria accounted for 25% of the 92% estimated malaria cases that occurred in Africa in 2017 where 217million malaria cases was reported (WHO, 2017) an indication of the endemicity of the country for malaria and other mosquito-borne diseases. The incidence of malaria, however, is reported to be increasing in Nigeria (WHO, 2018).

Invariably, Osun state is also endemic for mosquito-borne diseases (Adefioye *et al.*, 2007). Furthermore, the distribution of larval habitats and insecticide susceptibility status of *A. gambiae* complex is yet unknown in the State (Adeleke *et al.*, 2018). Thus, the need for the present study in assessing whether mosquitoes in Osogbo metropolis are resistant or susceptible to insecticides currently used for vector control of adult mosquitoes with a view to corroborating global efforts on the prevention and eradication of mosquito-borne diseases.

## Materials and Methods

### Study Site

The study was carried out in Osogbo, the capital city of Osun State, Nigeria. It is located on latitude (7° 49’N) and (7° 28’60’E) and at an elevation of 321m above the sea level in southwestern, Nigeria. The city experiences two seasons, the dry season (October to March) and wet season (April to September). The study was conducted between May and September 2022 during the wet season, which is associated with rainfall, thus the presence of breeding sites for the mosquitoes.

### Ethical Approval

Ethical approval was sought and obtained from the Osun State Ministry of Health, Abere, Osogbo, Osun State, Nigeria.

### Larval sampling and rearing to adult stage

Thorough quest was made to identify breeding sites for mosquito larval sampling. Larva collection was done using scoops, dippers, containers and sieves of about 0.55mm mesh size into well labelled transparent containers netted to prevent escape of those that may emerge to adult during transportation to the laboratory. In the laboratory, they are reared to adult stage under standard conditions before insecticide susceptibility testing. Adult mosquito identification was done under a dissecting microscope using morphological keys described by Hopkins (1953) and Gillet (1972).

### Bioassay

Emerged adult mosquitoes were tested for insecticide susceptibility/resistance using insecticide impregnated papers (4% DDT, 0.05% Deltamethrin, 0.75% Permethrin, Pirimiphos-methyl and OC-control) as described by the World Health Organization protocol (WHO, 2016). About 25 adult female mosquitoes were aspirated into each of the four exposure tubes for WHO bioassay containing insecticide impregnated papers of each of the insecticides used in an upright position to make four replicates for each insecticide and monitored for an hour. At the end of an hour, moribund (knockdown) mosquitoes were transferred to holding tubes while the dead ones are removed. A paper without insecticide was used as the control.

In the holding tubes, the moribund adult female mosquitoes are left for 24hrs at end of which the dead mosquitoes i.e., susceptible, and those that are alive i.e., resistant, were counted, recorded preserved using silica gel in an Eppendorf tubes. This process was repeated for adult female *A. gambiae* s. l., *A. aegypti* and *C. quinquefasciatus* mosquitoes respectively.

### Data Analysis

The data analysis was done using chi-square to test for the significant difference in insecticide resistance or susceptibility of the adult female mosquitoes to the insecticides used in the study, with a p-value of 0.05 indicating significance.

### Results

A total number of 1500 mosquito larvae were collected between May to September, 2022 in seven different locations within Osogbo metropolis. The mosquito larvae collected from different breeding sites are identified to belong to the genera: *Anopheles*, *Culex* and *Aedes* (Table 1).

**Table 1:** Mosquito larval genera composition in different habitats in the study area

| Habitat Types | Habitat frequency | Mosquito larval genera present |  |  | Percentage composition |
| --- | --- | --- | --- | --- | --- |
|  |  | *A. gambiae* s. l. | *C. quinquefasciatus* | *A. aegypti* | (%) |
| Gutter | 4 | 3 | 4 | 0 | 14.29 |
| Ground pool | 10 | 9 | 6 | 0 | 35.31 |
| Tyres | 4 | 0 | 0 | 4 | 14.29 |
| Abandon plastics | 2 | 0 | 0 | 2 | 7.14 |

After 60min of exposure, the highest kdr was observed with Pirimiphosmethy (85%) against *A. gambiae* s. l. while OC-control had lowest kdr (0%) against *A. aegypti* (p=0.69, p>0.005). It also recorded the highest susceptibility (69%) against *A. gambiae* s. l. after 24hours of exposure while OC-control had the lowest mortality rate (11%) against *A. aegypti* (p=0.65; p>0.05). Permethrin (49%) had highest resistance rate against *A. gambiae* s. l. over *C. quinquefasciatus* (24%) and *A. aegypti* (6%) while Pirimiphos-methyl showed the lowest resistance rate across the three mosquito genera (Table 2).

**Table 2:** kdr and mortality of the three mosquitoes species after 60min. and 24hrs exposures.

| Insecticide | Knockdown rate after 60min (%) |  |  | Female Mortality rate after 24hours (%) |  |  | Female Resistance after 24hours (%) |  |  |
| --- | --- | --- | --- | --- | --- | --- | --- | --- | --- |
|  | A. *gambiae* s. l. | C. *quinquefasciatus* | A. *aegypti* | A. *gambiae* s. l. | C. *quinquefasciatus* | A. *aegypti* | A. *aegypti* | C. *quinquefasciatus* | A. *aegypti* |
| Permethrin | 20 | 50 | 60 | 16 | 41 | 60 | 49 | 24 | 6 |
| Deltamethrin | 36 | 68 | 40 | 39 | 37 | 59 | 21 | 11 | 11 |
| Pirimiphos-methyl | 85 | 70 | 20 | 69 | 52 | 56 | 0 | 0 | 10 |
| DDT | 5 | 8 | 8 | 47 | 17 | 17 | 7 | 33 | 44 |
| OC-control | 1 | 0 | 0 | 26 | 15 | 11 | 20 | 38 | 45 |

Against *A. aegypti*, Permethrin had the highest kdr (60%) with OC-control having the lowest (0) after 60mins of exposure (Table 3). The *A*. *aegypti* had the highest resistance against OC-Control (45%)(p= 0.031; p< 0.05) (Figure 1). Permethrin had the highest susceptibility (60%) against *A. aegypti* while OC-control had the lowest (11%) (p= 0.005; p< 0.05) (Figure 1).

**Table 3:** kdr for *A. aegypti* after exposure for 60minutes.

| Insecticide (s) | Knockdown rate after 60min. |
| --- | --- |
| Permethrin | 60 |
| Deltamethrin | 40 |
| Pirimiphos-methyl | 20 |
| DDT | 8 |
| OC-control | 0 |

**Figure 1:**
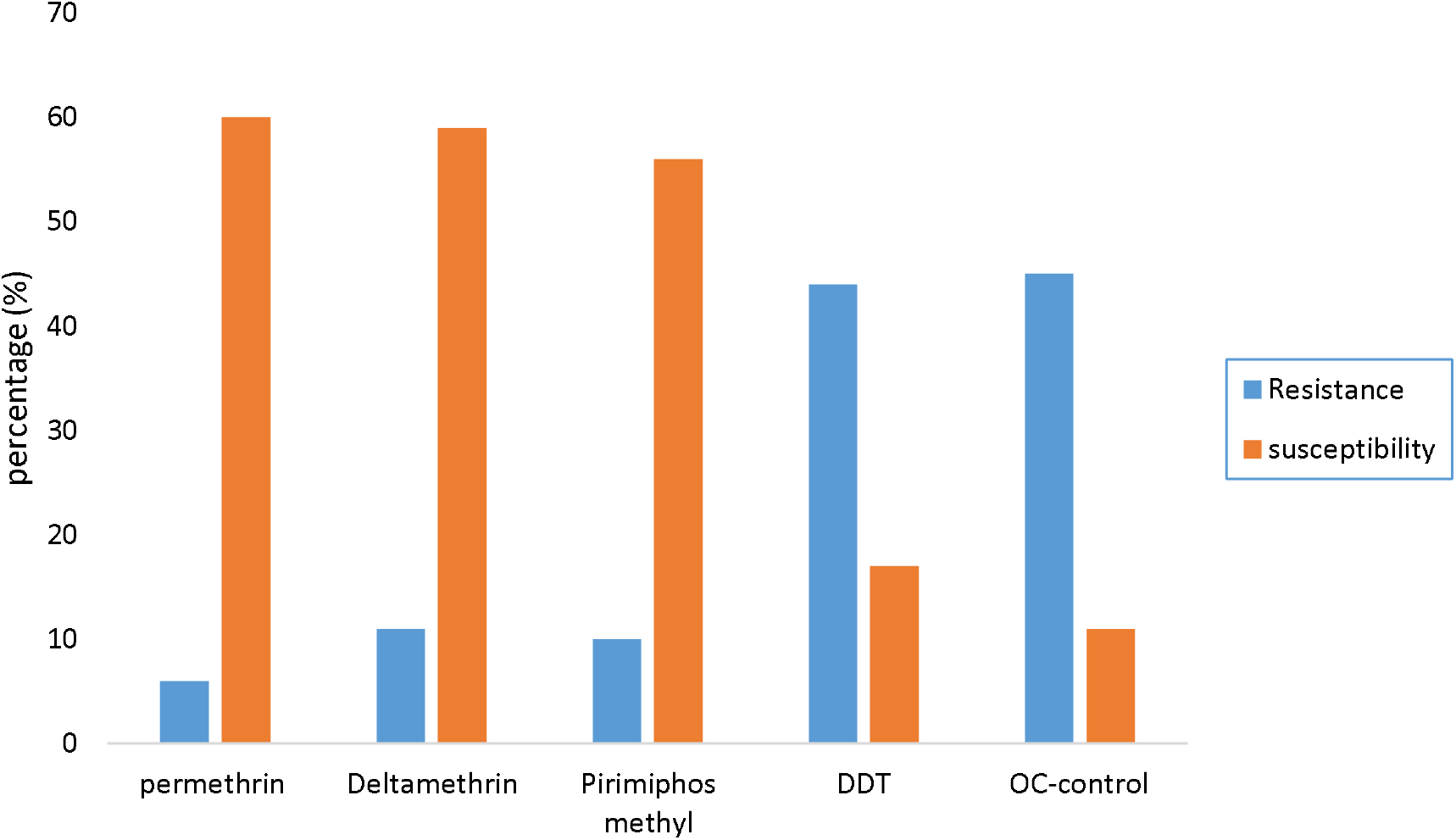
Resistance and susceptibility of *A. aegypti* after 24hrs exposure.

Pirimiphos-methyl had the highest kdr (70%) against *C. quinquefasciatus* while OC-control had the lowest (0%) after 60mins (Table 4). It had the greatest resistance to OC-control (38%), DDT (33%), Permethrin (24%), Deltamethrin (11%) and Pirimiphos-methyl (0%) respectively (p=0.016, p<0.05) (Figure 2). However, it was least susceptible to Pirimiphos-methyl (52%) and DDT (17%) respectively (p=0.013; p<0.05) (Figure 2).

**Table 4:** kdr for *C. quinquefasciatus* after 60min. exposure.

| Insecticide(s) | Kdr |
| --- | --- |
| Permethrin | 50 |
| Deltamethrin | 68 |
| Pirimiphos-methyl | 70 |
| DDT | 8 |
| OC-control | 0 |

**Figure 2:**
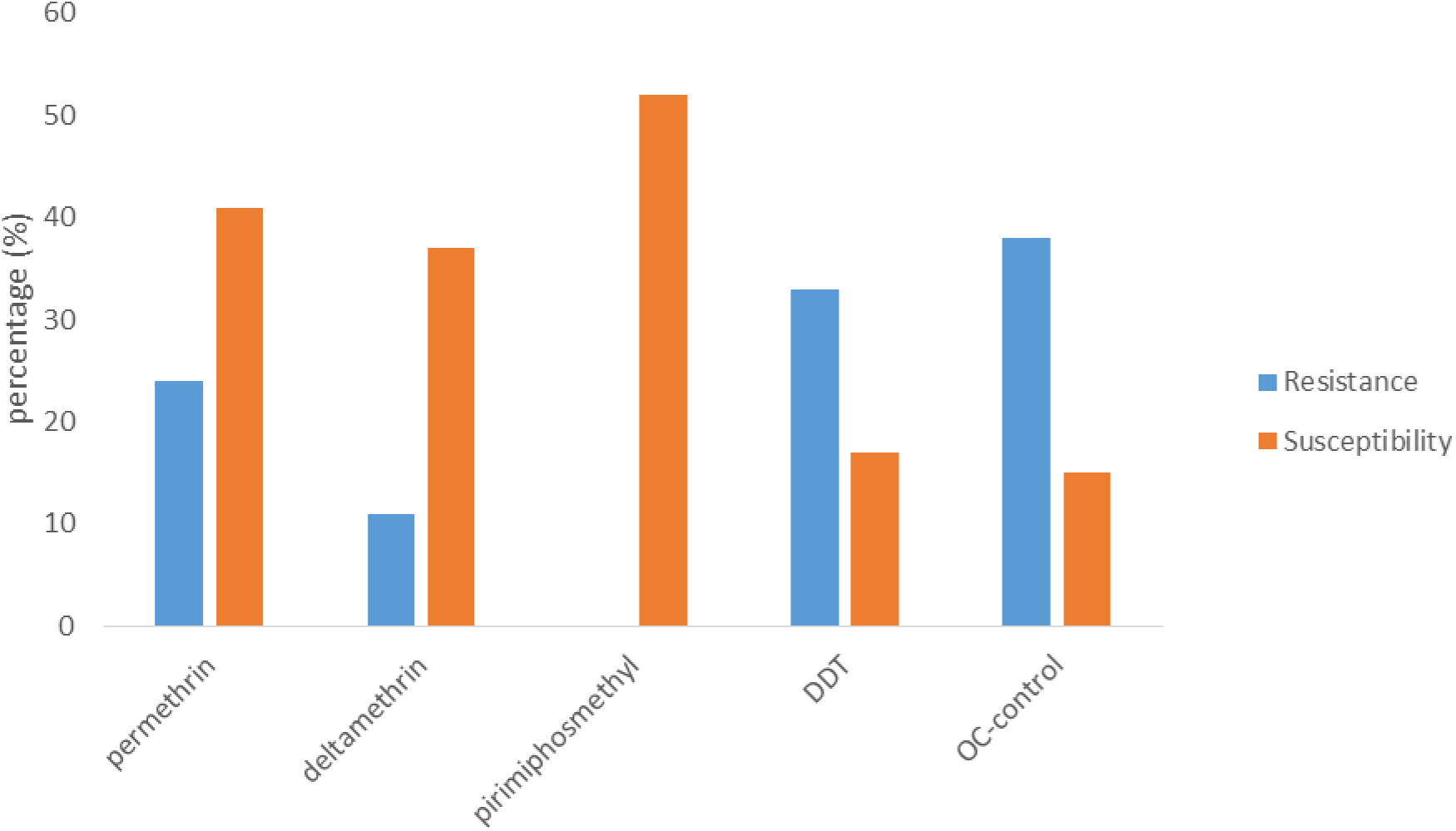
Resistance and susceptibility of *C. quinquefasciatus* after 24hrs exposure.

After 60min. exposure of *A*., Pirimiphosmethyl had the highest kdr (85%) with DDT having the lowest (5%)(Table 5). The vector showed the greatest resistance to Permethrin (49%), Deltamethrin (21%), OC-control (20%), DDT (5%) and Pirimiphos-methyl (0%) respectively (p=0.04, p<0.05) (Figure 3). Furthermore, the highest susceptibility was recorded with Pirimiphos-methyl (69) while the lowest was with Permethrin (16) (P=0.002; p<0.05) (Figure 3).

**Table 5:** **kdr of *A.****gambiae* s. l. **after 60min. exposure.**

| Insecticide(s) | Kdr |
| --- | --- |
| Permethrin | 20 |
| Deltamethrin | 36 |
| Pirimiphos-methyl | 85 |
| DDT | 5 |
| OC-control | 1 |

**Figure 3:**
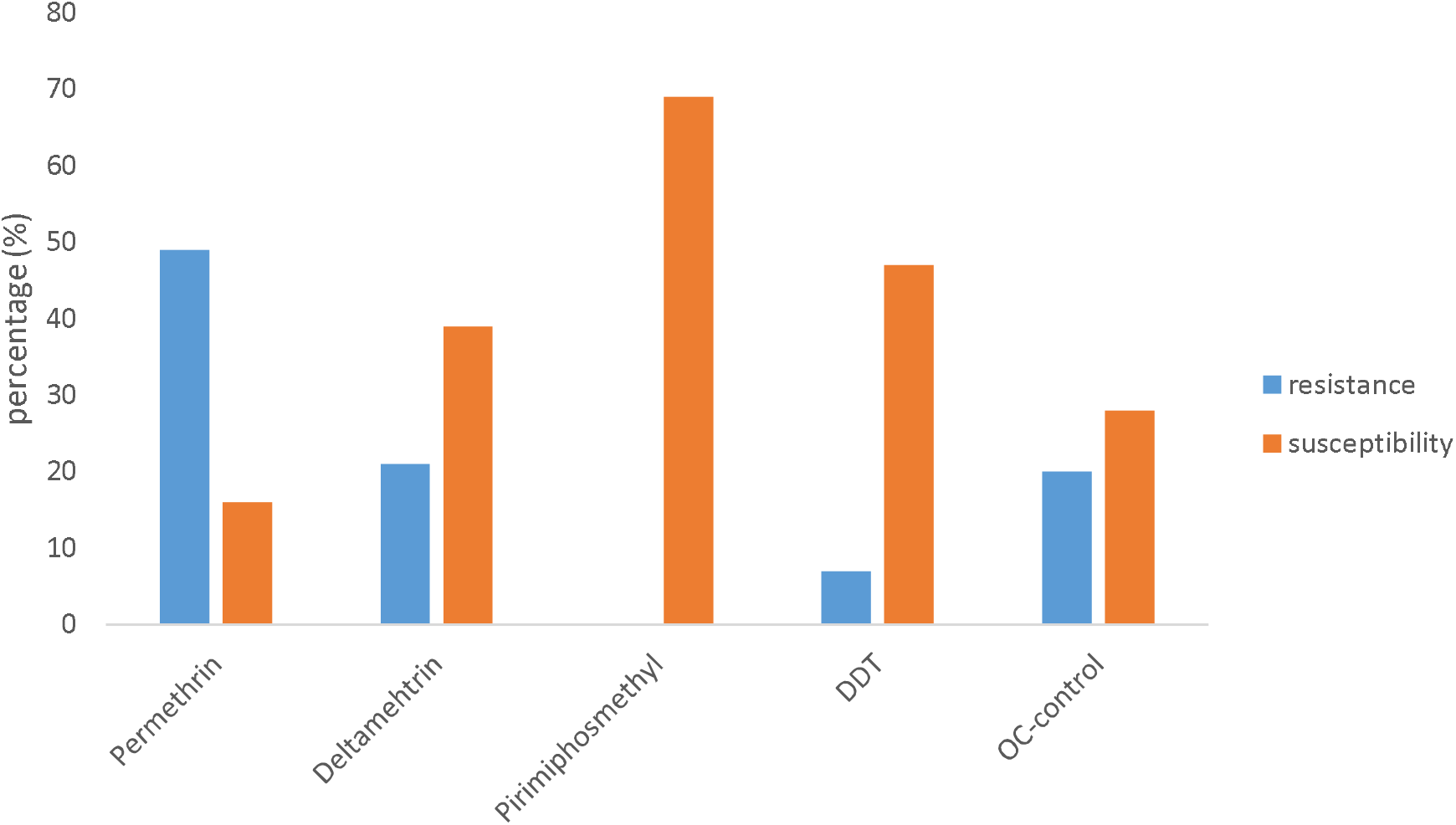
Resistasnce and susceptibility of *A. gambiae* s. l. after 24hrs exposure.

## Discussion

Knock down rate of the vectors showed that the kdr increases as the period of exposure increases. During the kdr assessment for the mosquito vectors the highest kdr was recorded in Pirimiphos-methyl against the three species after exposure (p=0.04; p<0.05). This is followed by Permethrin (p=0.02; p<0.05), Deltamethrin (p=0.008; p>0.05), DDT and OC-control having the lowest kdr (p=0.04; p<0.05). The assessment of the kdr showed that the tested impregnated paper induced knockdown of the adult female mosquito vectors, suggesting that knockdown mechanism could be operating in the local mosquito populations in the study area. This conforms with earlier studies which indicates the knockdown effect of impregnated papers against the mosquito species in Nigeria (Oduola *et al.*, 2010; Olayemi *et al.*, 2011; Umar *et al.*, 2014; Awolola *et al.*, 2018)

After 24hrs exposure, *A. gambiae* s. l.*, C. quinquefasciatus* and *A. aegypti* were resistance to OC-control (p=0.009; p>0.05). This could be due to reported cases of vector resistance to organochlorine class of insecticides (Adeogun *et al.*, 2022). According to the WHO (1996), *Culex, Aedes* and *Anopheles* species have developed resistance rapidly to various insecticides in many countries, but *Culex* can develop resistance more rapidly to insecticides than other mosquitoes. The high resistance of *A. gambiae* s. l.*, C. quinquefasciatus* and *A. aegypti* compared with OC-control and DDT (p=0.0004; p<0.05), is in conformity with previous studies where emerging resistance to DDT have been reported (Adeleke *et al.*, 2018). Resistance was high but moderate in Permethrin (48%) and Deltamethrin (60%) (p=0.002; p<0.05). The vectors showed a high susceptibility to Pirimiphos-methyl (70%) (p=0.002; p<0.05) and in consonance with findings by (Corbel *et al.*, 2007). Nkya *et al.*, (2014) reported that the mosquito species were susceptible to organophosphate but resistance to DDT with reduced susceptibility to Pyrethroid (Permethrin and Deltamethrin).

The low mortality rate with OC-control and DDT could be because of the presence of kdr mutation which is conferring resistance to the vectors since the spade of rapid resistance to insecticide by the vectors has been attributed to mutation. This could also be because of environmental factors (Nkya *et al.*, 2014) since environmental management is germane in vector control and disease elimination.

## Conclusion

Owing to the endemicity of mosquito-borne diseases such as malaria, lymphatic filariasis etc. in the state and globally, the susceptibility of *A. gambiae* s. l. and *C. quinquefasciatus* to Pirimiphos-methyl and *A. aegypti* to Permethrin respectively. According to the present study, we suggest the possibility of success of vector control using Pirimiphos-methyl against *A. gambiae* s. l. and *C. quinquefasciatus* and Permethrin against *A. aegypti.* Thus, the need for the state government and health agencies to employ these insecticides in mosquito-borne diseases vector control programmes in the state.

